# Redox Buffering Capacity of Nanomaterials as an Index of ROS-based Therapeutics and Toxicity: A Preclinical Animal Study

**DOI:** 10.1101/2021.03.14.435286

**Authors:** Aniruddha Adhikari, Susmita Mondal, Monojit Das, Ria Ghosh, Pritam Biswas, Soumendra Darbar, Soumendra Singh, Anjan Kumar Das, Siddhartha Sankar Bhattacharya, Debasish Pal, Asim Kumar Mallick, Samir Kumar Pal

## Abstract

Precise control of intracellular redox status, i.e., maintenance of physiological level of reactive oxygen species (ROS) for mediating normal cellular functions (oxidative eustress) while evading the excess ROS stress (distress) is central to the concept of redox medicine. In this regard, engineered nanoparticles with unique ROS generation, transition, or depletion functions have the potential to be the choice of redox therapeutics. However, it is always challenging to estimate whether ROS-induced intracellular events are beneficial or deleterious to the cell. Here, we propose the concept of redox buffering capacity as a therapeutic index of engineered nanomaterials. As a steady redox state is maintained for normal functioning cells, we hypothesize that the ability of a nanomaterial to preserve this homeostatic condition will dictate its therapeutic efficacy. Additionally, the redox buffering capacity is expected to provide information about the nanoparticle toxicity. Here, using citrate functionalized trimanganese tetroxide nanoparticles (C-Mn_3_O_4_ NPs) as a model nanosystem we explored its redox buffering capacity in erythrocytes. Furthermore, we went on to study the chronic toxic effect (if any) of this nanomaterial in animal model in order to co-relate with the experimentally estimated redox buffering capacity. This study could function as a framework for assessing the capability of a nanomaterial as redox medicine (whether maintains eustress or damages by creating distress), thus orienting its application and safety for clinical use.

## INTRODUCTION

Reactive oxygen species (ROS) and associated oxidative stress (so-called) remains the most fascinating paradigm for elucidating the adverse effects of engineered nanomaterials at cellular and molecular level [1]. ROS generating nanomaterials are generally considered cytotoxic for biomedical use, and their therapeutic applications are restricted to anticancer or anti-microbial activities [2-5]. Indeed, elevated ROS levels have the capacity to oxidize unsaturated fatty acids in lipids, and amino acids in proteins, inducing irreversible damage to vital organelles, and DNA, ultimately leading to cell apoptosis, and necrosis resulting in various chronic disorders including neurodegenerative, diabetes, and cardiovascular diseases [6-8]. However, recent understandings in free radical biology illustrate the pivotal role of ROS (at low concentration) as second messengers to modulate essential cellular functions like signaling, adhesion, migration, proliferation and homeostasis [9]. This inherent duality of ROS, the purposeful beneficial functions at physiological level (i.e., oxidative eustress), as well as, deleterious effects at supraphysiological concentration (i.e., oxidative distress) highlights the importance of maintaining proper redox balance for healthy organismal functions, and leads to the concept of redox medicine [9, 10]. Redox medicine epitomize a promising tool for restoration of cell and tissue homeostasis via direct detoxification of reactive intermediates and/or by triggering cytoprotective and antioxidant signaling pathways [10]. In this sense, an interesting (perhaps the most important) avenue in redox medicine is the maintenance of oxidative eustress to regulate redox signaling and associated cell functions in addition to modulation of oxidative distress (i.e., inhibition to counteract pathological cell loss and induction to promote cell death of pathological cells). The disappointing history of efforts to prevent, or treat diseases with exogenous antioxidants (e.g., vitamin c, ascorbic acid, polyphenols etc.) further reflect the importance of redox-based therapeutics for clinical translation [11].

In this new front of redox medicine, ROS-based nanomedicines that involve nanomaterials with ROS-regulating properties, holds promise for optimized therapeutic efficacies [9]. Recent advancement in controlled synthesis process resulted into a wave of multifunctional nanomaterials with unique ROS generation, transition, or depletion functions. However, for *in vivo* application, it is a double-edged sword. After exerting expected therapeutic functions, the elevated ROS level may also impair adjacent normal tissues [5]. Moreover, if the nanomaterials are not degraded or excreted, their ROS-generating functions may still exist that may result in sustained oxidative damage. Therefore, balancing the therapeutic outcome and side effect of ROS generating nanomaterials is important for the optimization of therapeutic effects, which needs more careful characterization of the *in vivo* behavior of ROS and more exhaustive safety assessment of the nanomaterials [9]. In other words, information about ROS-mediated changes in cellular redox state, or ability of a nanomaterial to maintain eustress condition could be instrumental in dictating the therapeutic potential of a nanomaterial.

Unfortunately, the quantitative basis of changes in the intracellular redox state of the cells is not well-defined, thus leading to the dilemma that redox changes induced by oxidants in distinct cells types cannot be predicted [12]. The current state of the art measurement methods for estimating changes in cellular redox state is by measuring the concentration variation of intracellular redox couple GSSG/2GSH, and calculating the reduction potentials using Nernst equation, which is grossly misleading as spatial-temporal control exists between redox couples, and information about single couple cannot reflect the actual redox status [12-14]. Moreover, the method is cumbersome and prone to errors.

In this study, we introduce the concept of redox buffering capacity as a therapeutic index of engineered nanomaterials. As a steady redox state is maintained for normal functioning cells, we hypothesize that the ability of a nanomaterial to preserve this homeostatic condition will dictate its therapeutic efficacy. Additionally, the redox buffering capacity is expected to provide information about the nanoparticle toxicity. Here, using citrate functionalized trimanganese tetroxide nanoparticles (C-Mn_3_O_4_ NPs) as a model nanosystem we explored its redox buffering capacity in erythrocytes. In this regard it is worth mentioning that, C-Mn_3_O_4_ NPs has previously been well studied from our group and showed therapeutic efficacy using both ROS generation and ROS scavenging abilities in *in vitro* as well as *in vivo* models. It treats hyperbilirubinemia by oxidative breakdown of bilirubin inside the body. On the other hand, it can mimic glutathione peroxidase to scavenge ROS and treat several diseases including hepatic fibrosis, heavy metal toxicity and Huntington’s disorder. Thus, it is of considerable interest to assess the redox modulatory properties of this multifunctional nanoparticle.

Furthermore, we went on to study the chronic toxic effect (if any) of this nanomaterial in animal model in order to co-relate with the experimentally estimated redox buffering capacity. This study could function as a framework for assessing the capability of a nanomaterial as redox medicine (whether maintains eustress or damages by creating distress), thus orienting its application and safety for clinical use.

## RESULTS AND DISCUSSION

### Redox Buffering Capacity of C-Mn_3_O_4_ NPs

In the present study we used ∼10 nm nearly spherical C-Mn_3_O_4_ NPs having monomodal size distribution (Figure 1). The optical properties of the nanoparticles have been thoroughly investigated and described by us in previous studies [15, 16]. The nanoparticle has the unique ability to generate ROS in dark, without any photo- or chemi-activation [17]. Moreover, the NPs can function as a GSH-dependent GPx mimic to scavenge H_2_O_2_ in physiological *milieu*.

**Figure 1.**
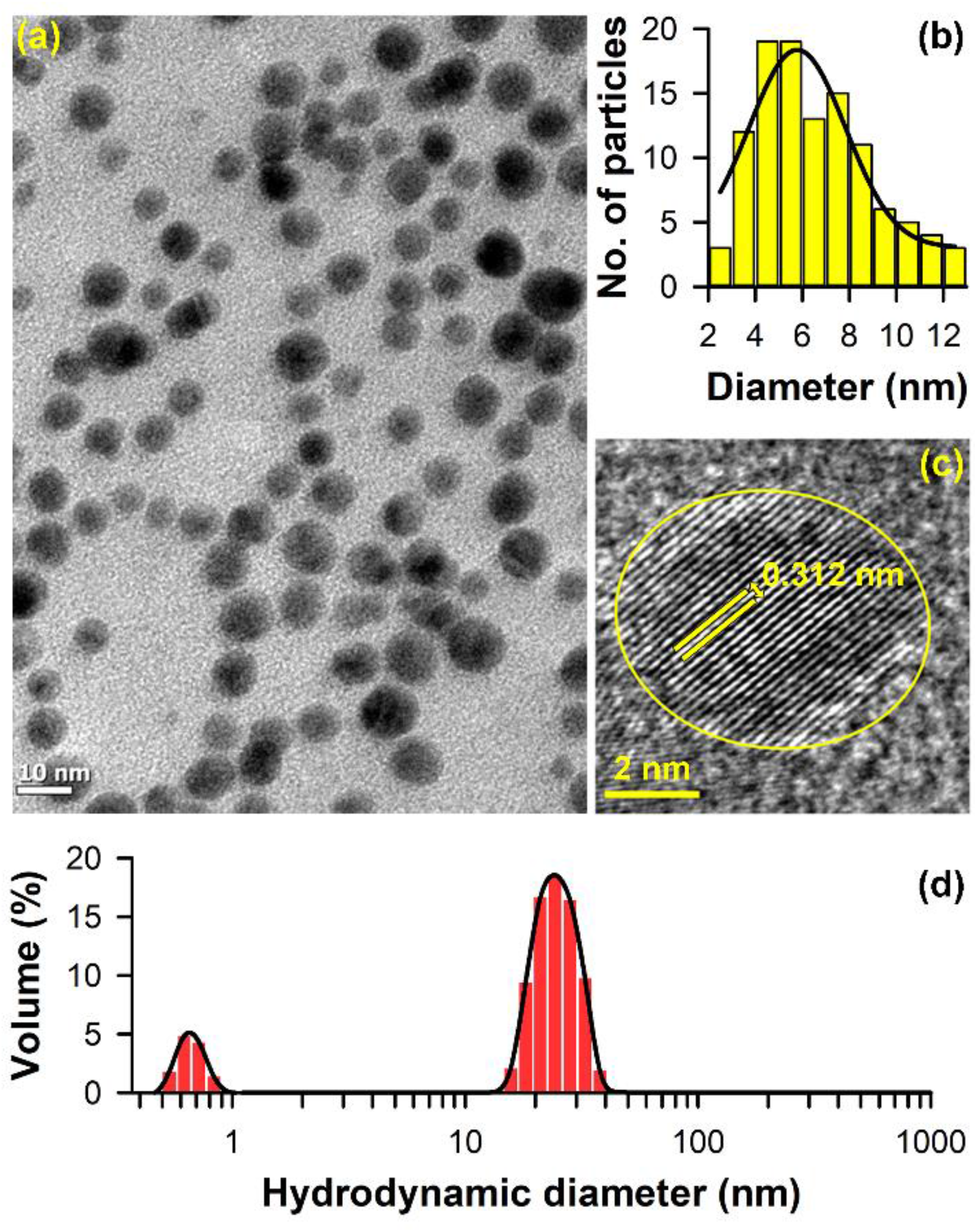
Properties of citrate-Mn_3_O_4_ nanoparticles (C-Mn_3_O_4_ NPs). (a) Transmission electron microscopic (TEM) images of C-Mn_3_O_4_ NPs. (b) The particle size distribution of the NPs with average size of 6.25±2.41 nm. (c) High resolution TEM (HR-TEM) of single NP shows the interplanar distance of 0.312 nm for 112 planes of C-Mn_3_O_4_ NPs. (d) Hydrodynamic diameter of C-Mn_3_O_4_ NPs as measured by DLS.

In order to explore the redox buffering effect of C-Mn_3_O_4_ NPs, we used the DCFH assay and measured the DCF intensity as a quantitative marker of ROS present in the reaction medium. First, we evaluated the *in vitro* redox buffering capacity of the synthesized NPs using H_2_O_2_ as an oxidant molecule. Figure 2a reflects the rate of DCFH oxidation with an increasing concentrations of H_2_O_2_ added to aqueous medium. The results clearly shows that the DCFH oxidation increased significantly with increasing concentration of H_2_O_2_. On the other hand, introduction of C-Mn_3_O_4_ NPs into the medium resulted into insignificant alteration in DCF intensity even after the addition of the highest concentration of H_2_O_2_. As shown in Figure 2a, the DCFH oxidation rate remained almost constant up to the addition of 2.5 mM of H_2_O_2_. It signifies the efficient quenching of ROS generated from H_2_O_2_ by C-Mn_3_O_4_ NPs. It should be noted that C-Mn_3_O_4_ NPs always maintained a constant ROS concentration in the reaction medium instead of eliminating all the ROS. Figure 2a depicts the relation between the DCFH oxidation rate (V_f_) and oxidant (H_2_O_2_) concentration. Here, it is evident that DCFH oxidation by H_2_O_2_ follows an exponential kinetics and the oxidation rate clearly reduced in presence of C-Mn_3_O_4_ NPs.

**Figure 2.**
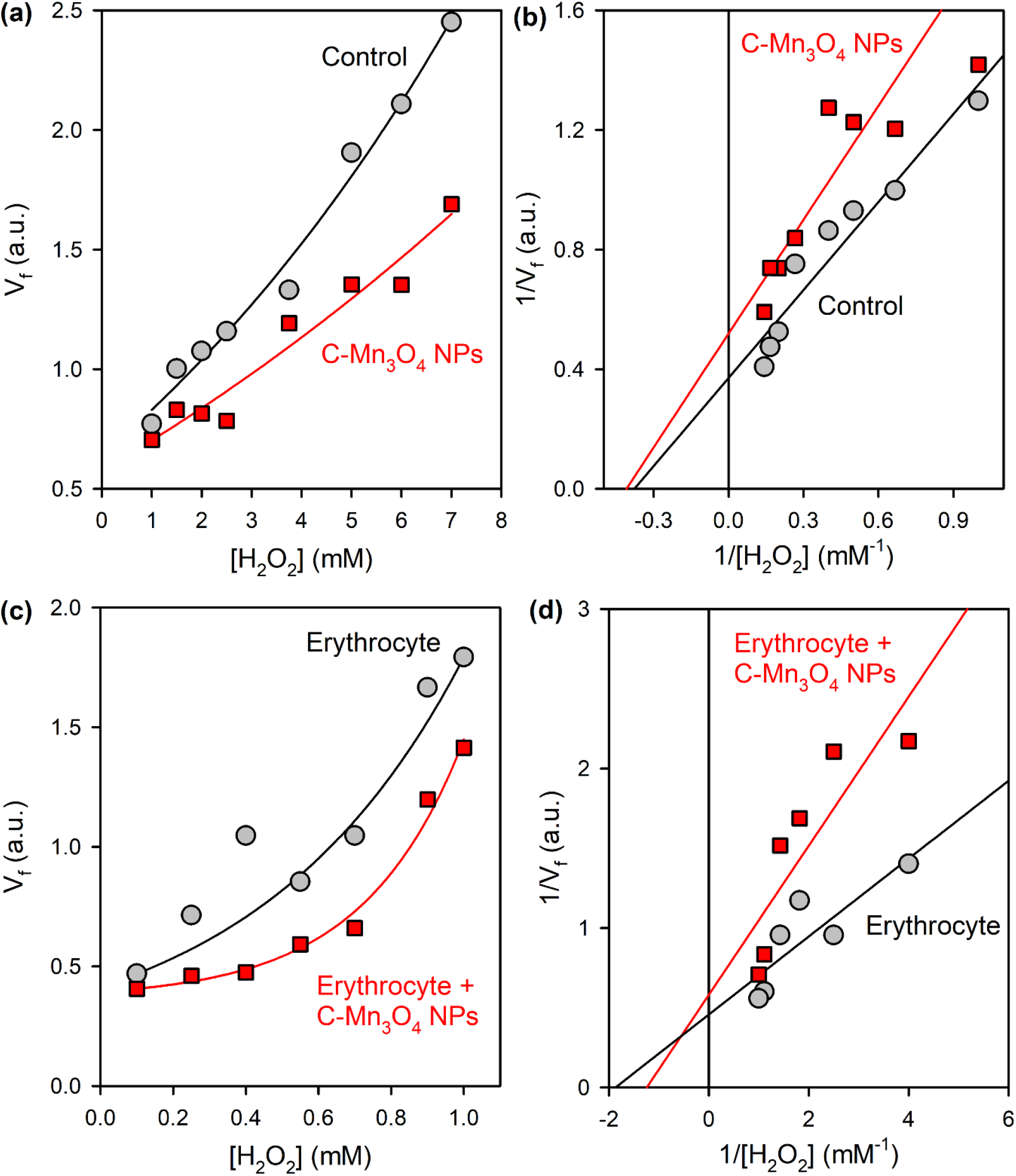
Redox buffering capacity of C-Mn_3_O_4_ NPs. (a) Rate of DCFH oxidation in presence of H_2_O_2_. (b) Double reciprocal plot of the same. (c) Rate of DCFH oxidation in presence of H_2_O_2_ in erythrocytes. (d) Double reciprocal plot of the same.

Next we quantified the *in-vitro* redox buffering capacity of the synthesized NPs. As per definition, the redox buffer capacity of a system is numerically equal to the magnitude of change of concentration of an oxidant (or a reductant) added to a solution, which is reduced or oxidized, to change the effective reduction potential by 1 unit (1 V) [12]. Thus, the redox buffer capacity (β) can be represented as:

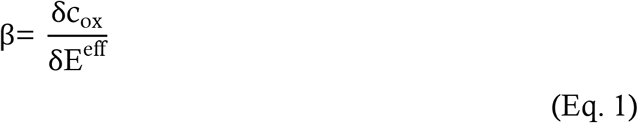

Where c_ox_ represents the concentration of the oxidant added to the system.

Experimentally, the redox buffering capacity and the effective reduction potential can be determined from the dependence of 1/V_f_ on 1/c_ox_ (Figure 2b). The correlation can be described by the following equation:

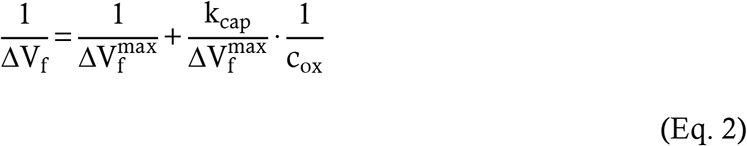

Where, 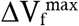 is the highest rate of DCFH oxidation and k_cap_ is a constant having dimension of concentration. According to Eq. 2, the redox buffering capacity 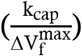 can be calculated from the tangent of slope angle of the plot of 1/V_f_ (Y-axis) vs. 1/c_ox_ (X-axis).

Figure 2b shows the significantly higher (∼1.3 times) redox buffering capacity of C-Mn_3_O_4_ NPs (1.29 unit) compared to Mili-Q water (0.97 unit).

To test the aforementioned redox buffering ability of C-Mn_3_O_4_ NPs in the cellular *milieu* we used human erythrocytes, or red blood corpuscle (RBC). The choice of erythrocytes was based on the fact that (i) their continuous exposure to high oxygen tensions, (ii) inability to replace damaged components, and (iii) the composition of membrane lipids partly of polyunsaturated fatty acid side chains that are vulnerable to peroxidation. Exogenously introduced H_2_O_2_ served as the inducer of oxidative stress. It is well-known that H_2_O_2_ generates massive oxidative stress and perturbs the intracellular redox state. For living cells, the electron donating and electron accepting capabilities of the intracellular medium can be specified by the value of effective redox potential, and any change in effective redox potential can be measured using specific redox sensor like DCFH in our study. Figure 2c and 2d represent the *in-cellulo* redox buffering capacity of C-Mn_3_O_4_ NPs. It is evident from the results that the DCF intensity of the RBCs treated with C-Mn_3_O_4_ NPs is significantly lower than the control RBCs treated with normal saline (Figure 2c). Following the method described above we found that the redox buffering capacity of C-Mn_3_O_4_ NPs (0.46 units) when integrated in erythrocytes is significantly higher (∼1.9 times) compared to the intrinsic cellular components (0.24 units) (Figure 2d). It has to be noted that the redox buffering capacity of the C-Mn_3_O_4_ NPs in cellular system is considerably higher compared to *in vitro* aqueous system. This may be due to the additional (i.e., other than direct antioxidant activity) GPx mimicking activity of the C-Mn_3_O_4_ NPs [18] which can only happen in presence of GSH, and intracellular metabolite.

Redox buffering requires sensing and rapid adjustment in the redox environment to maintain the oxidative eustress in cells. Cumulative data obtained from the experiments (both *in-vitro* and *in-cellulo*) suggest that C-Mn_3_O_4_ NPs can sense and swiftly shift the redox state in the favour of oxidative eustress. This behaviour of C-Mn_3_O_4_ NPs as a redox modulator is consistent with that of pH buffers. pH buffers adjust the pH of the solution by sensing the concentration of H^+^ ions present. Similarly, C-Mn_3_O_4_ NPs perceive the amount of ROS in the medium and change its redox state accordingly. Also, it has been shown that a significant change in the concentration of C-Mn_3_O_4_ NPs in the solution can cause a change in the redox state just like a pH buffer.

Previous studies suggest that for maintaining the oxidative eustress, C-Mn_3_O_4_ NPs generate ROS from dissolved O_2_ and H_2_O through continuous disproportionation and comproportionation of the surface Mn^2+^/Mn^3+^/Mn^4+^ ions present at the spinel structure [15]. Electrons, produced from this process of conversion of Mn^2+^→ Mn^3+^→ Mn^4+^, react with the dissolved O_2_ and H_2_O of the medium and produce ROS. The ROS produced in this process is simultaneously neutralized by the conversion of Mn^4+^→ Mn^3+^→ Mn^2+^, i.e. the opposite pathway [17, 19, 20]. Therefore, when the cells stay at oxidative eustress condition, a dynamic equilibrium is maintained between the generation and elimination of ROS by C-Mn_3_O_4_ NPs. However, in many instances, the ROS level in the cells decreases resulting into immune suppression. During this condition the ROS generated by C-Mn_3_O_4_ NPs can take part in the cellular signalling process and maintain the normal cell functions. On the other hand, in oxidative distress conditions, the scenario becomes more complicated. The C-Mn_3_O_4_ NPs can follow numerous buffering strategies to mitigate the oxidative distress. Besides quenching of ROS through the conversion of Mn^2+^/Mn^3+^/Mn^4+^, another likely strategy is that, in ROS excess medium, NPs act as a cofactor to some antioxidant enzymes and directly help intracellular anti-oxidant defence system [17, 20, 21]. Furthermore, the NPs can function as a GSH-dependent GPx mimic and scavenge excess ROS being incorporated into the cellular antioxidant enzyme network, as described in our previous study.

The combined results suggest that the C-Mn_3_O_4_ NPs can efficiently function as a redox buffer in intracellular condition. The described ‘redox titration’ method could be beneficial for comparing buffering capacity, as a marker of cyto-compatibility of different nanomaterials using minimum resources.

### C-Mn_3_O_4_ NPs show excellent hemocompatibility

Next we evaluated chronic toxicity of C-Mn_3_O_4_ NPs in animal model. With all newly developed nanoparticles, the potential for toxicity remains a factor in determining their specific applications in experimental small-animal, and ultimately clinical application. When NPs are administered *in vivo*, the most important physiological system they interact with is the blood and blood components. Then, if cleared by immune cells, they leave away from the bloodstream and reach the target organ [22]. Thus, it is important to measure the hemocompatibility of the NPs targeted for therapeutic use. We performed the hemolysis assay for evaluation of the hemolytic behaviors of C-Mn_3_O_4_ NPs on isolated mouse erythrocytes. The erythrocytes were first isolated by centrifugation and purified by sequential washes with sterile isotonic phosphate-buffered saline (PBS) pH 7.4, then diluted to 5% hematocrit and incubated with different concentrations of NP suspensions (0.01, 0.05- or 0.1-ml ml^-1^ PBS). The same amount of PBS was used as the negative control. An aqueous solution of RBCs was used as a positive control that is likely to show hemolysis. After 6 hrs incubation at room temperature, the samples were spun down to check the extent of hemolysis. Considering the hemolytic ability of positive control (water) as 100%, all the NP treated samples showed negligible (<5%) hemolysis (Figure 3a and 3b). Corresponding absorbance spectra (with the supernatant after final centrifugation) showed the optical features of C-Mn_3_O_4_ NPs (Figure 3c) with ligand to metal charge transfer band at 300 nm and d-d transition bands at 429, 526 nm. Only the positive control sample showed a distinct absorbance signature of hemoglobin with the Sorret band at 420 nm and Q-bands at 546 and 576 nm (Figure 3c). The scanning electron micrographs (SEM) (Figure 3e) of erythrocytes and Leishman stained whole blood smears supported the same (Figure 3f). The cell membrane remained unaltered after particle interaction, and the erythrocytes maintained normal biconcave shape just like control ones (Figure 3e & 3f). To study the *in vivo* effect of C-Mn_3_O_4_ NPs on blood, different hematology and serum biochemical indicators were measured after 90-days administration of the particles. All hematology marker values were within normal ranges and did not indicate a trend of toxicity associated with NPs (Supplementary Table S1). We further went on to assay ALAD activity, an important enzyme in the hematopoietic pathway which usually gets downregulated in cases of metal toxicity, particularly due to exposure of heavy metals like lead, mercury, etc. [23] However, we found no change in the activity of ALAD (Figure 3d). The hematoxylin and eosin stained spleen showed no abnormality (Figure 3g). These results indicate C-Mn_3_O_4_ NPs have excellent hemocompatibility.

**Figure 3.**
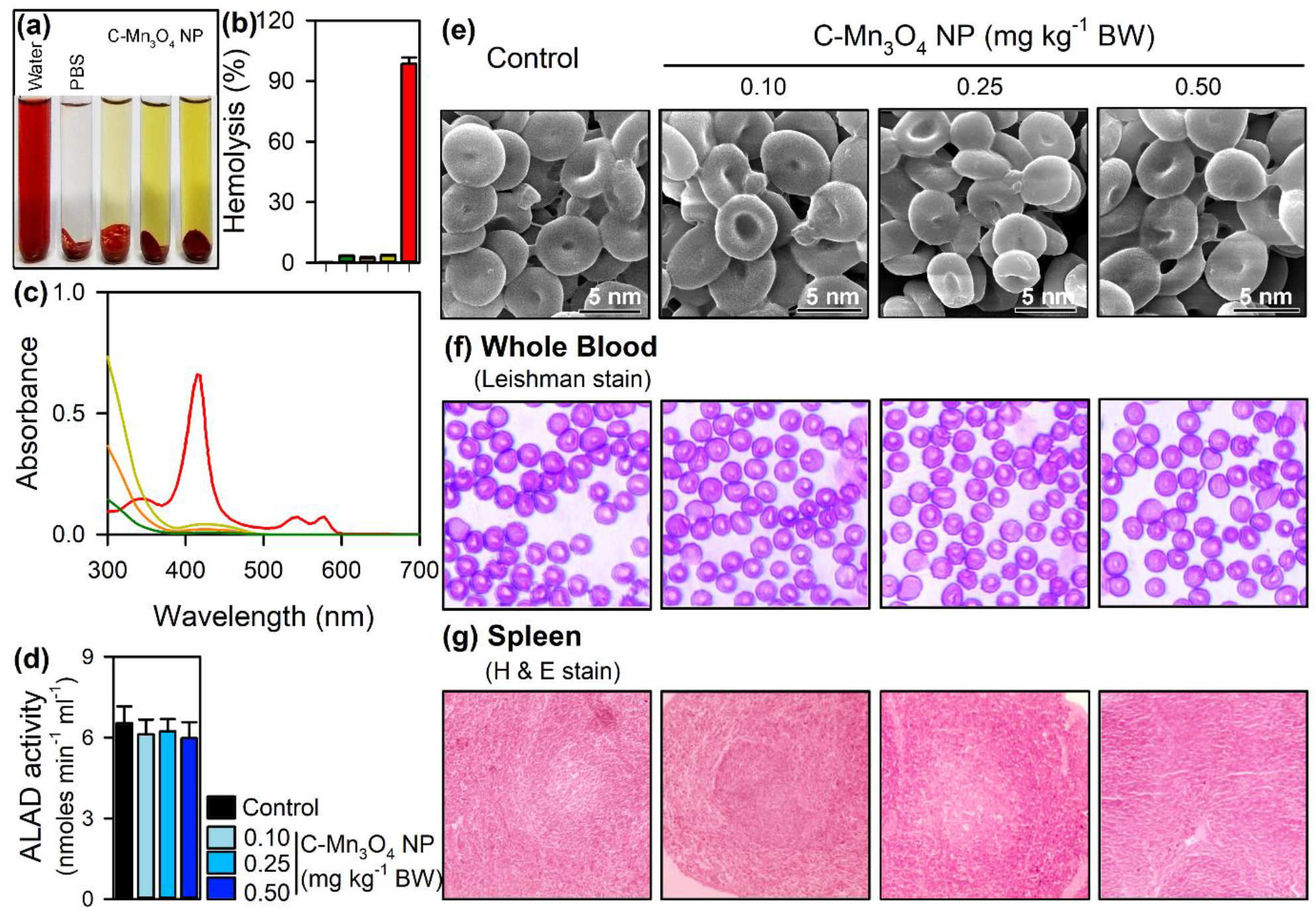
Effect of C-Mn_3_O_4_ NPs on haematological system. (a) Visual inspection of the tubes containing RBC after exposure to NPs. No visible haemolytic behaviour was observed for NPs, water was used as positive control and PBs as negative control. The bar plot shows percentage of haemolysis induced by C-Mn_3_O_4_ NPs. (c) UV-vis absorbance spectra of the supernatant. (d) ALAD activity. (e) Scanning electron micrographs of the RBCs incubated with C-Mn_3_O_4_ NPs shows no visible damage to the membrane or morphology. (f) Leishman stained whole blood from the BALB/c mice treated with different doses of NPs for 90 days. (g) H&E stained spleen sections after 90 days’ oral administration of NPs. All data represented as Mean ± Standard Deviation (SD). N=6 for each measurement.

### High dose of C-Mn_3_O_4_ NPs cause mild oxidative damage to liver

The results of biodistribution (Supplementary Figure S1) study suggest the affinity of C-Mn_3_O_4_ NPs towards the liver. The gross morphology (Figure 4a) and relative liver weights do not reveal any toxic effect on the organ. But compared to control animals, 0.5 mg kg^-1^ BW NP-treated mice (not the other two groups of NP treated ones) displayed significantly higher serum alanine aminotransferase (ALT) and aspartate aminotransferase (AST) activity (Figure 4b & 4c, p<0.001 in one way ANOVA). A similar trend was observed in the case of alkaline phosphatase (ALP) and γ-glutamyl transferase (γGT) activity (Figure 4d & 4e), however, bilirubin levels showed no significant differences among groups (Figure 4f). The increased serum activity of transferases, normally confined within hepatic cells (not in serum), can be attributed to the damaged structural integrity of hepatocytes and subsequent release into the circulation. To know the real reason behind such elevated liver function parameters in the case of 0.5 mg kg^-1^ BW NP-treated mice we went of histological analysis. In vehicle control animals, liver sections showed normal hepatic cells with well-preserved cytoplasm, prominent nucleus and nucleolus, and central vein (Figure 4a). They along with 0.1 and 0.25 mg kg^-1^ BW NP-treated livers displayed characteristic hepatic architecture with hepatic plates directed from the portal triads toward the central vein where they freely anastomose. Inflammatory cells were seen intervening the hepatic strands and surrounding degenerative hepatocytes (Figure 4a). In addition, pericentral vein inflammation was also seen. Further histological damages including necrosis, ballooning degeneration, vacuolation, increased mitosis, and calcification were observed in the same group of mice (Figure 4a). Thus, it is evident from the results of histopathological observations and liver function tests that C-Mn_3_O_4_ NPs at the highest administered dose (0.5 mg kg^-1^ BW) can cause moderate hepatic toxicity. Previously, we have demonstrated the NPs ability to generate reactive oxygen species (ROS) in aqueous solution. So, to understand the pathophysiological mechanism of observed toxicity, we measured plasma thiobarbituric acid reactive substances (TBARS) or malondialdehyde (MDA) content, a byproduct of lipid peroxidation and also an index of oxidative damage, which was found to be high in 0.5 mg kg^-1^ BW NP-treated group (Figure 4i). We further extended our investigation to the changes in activities of superoxide dismutase (SOD), catalase (CAT) and glutathione peroxidase (GPX) which creates a mutually supportive team to fight against increased ROS [17, 24]. We found the downregulation of all the antioxidant enzyme activities in 0.5 mg kg^-1^ BW NP-treated mice compared to control (Figure 4j-4l). The mitochondrial markers of oxidative damage like mPTP opening, membrane potential, and ATP content followed same trend (Figure 4m-4o). Though previous studies have shown that, Mn_3_O_4_ NPs can display antioxidant enzyme mimetic (particularly SOD and CAT) activity in the cellular environment, the humongous amount of ROS instantaneously generated due to high local concentration of NPs cannot be quenched through that mechanism. It should be noted that at the therapeutic dose (0.25 mg kg^-1^ BW NPs) there is no evidence of oxidative damage.

**Figure 4.**
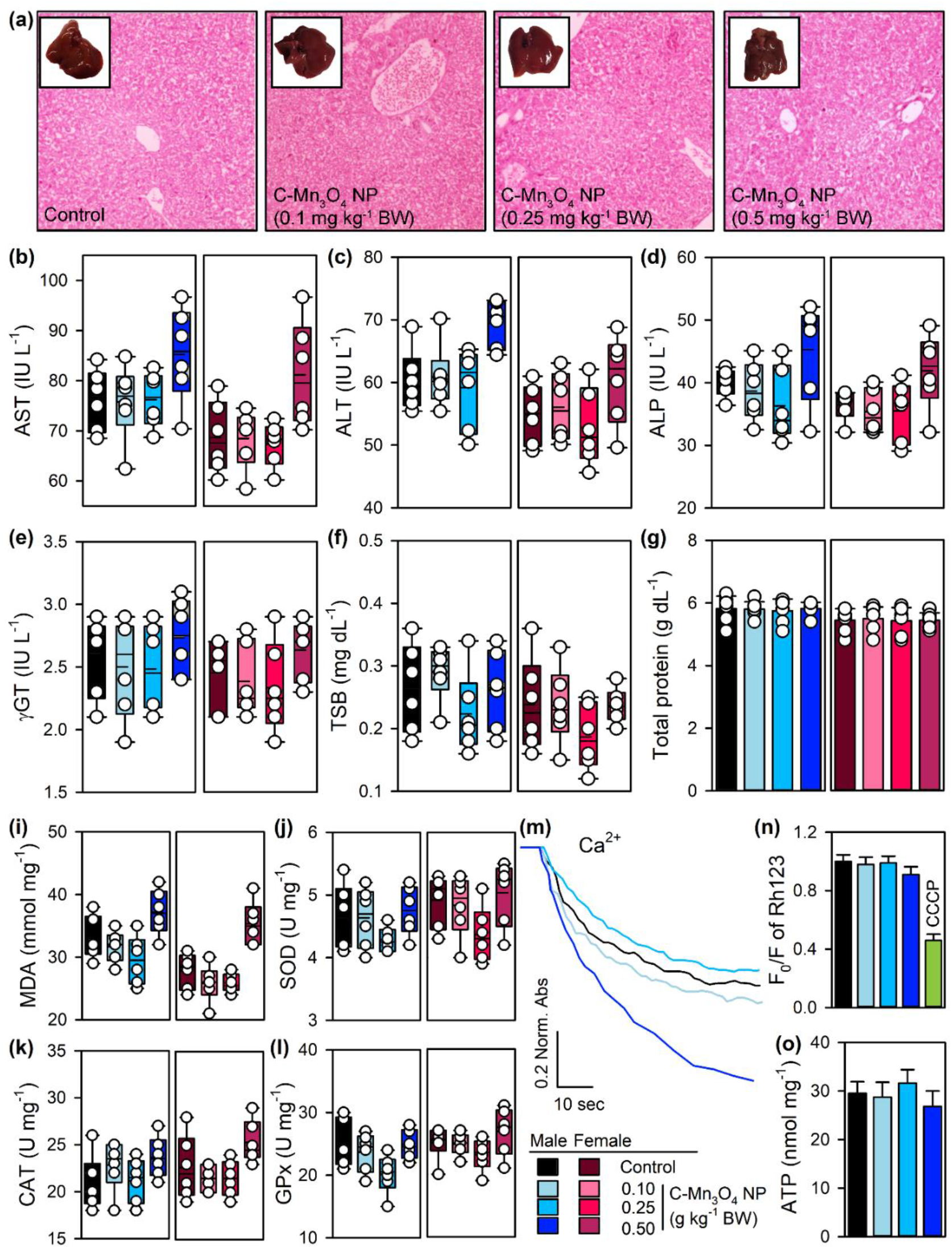
Effect of C-Mn_3_O_4_ NPs on hepatic function after chronic administration. (a) Morphology of the liver and H&E stained liver sections. (b-g) Effect on liver function parameters show chronic administration of high dose of NPs can cause damage hepatic cells. However, the therapeutic dose has no detrimental effect. (i-l) Assessment of cellular antioxidant enzymes revealed the damage to be ROS mediated. (m-o) Effect of the NPs on mitochondria reveals membrane depolarization along with increased MPTP which in turn increase cellular apoptosis through release of cytochrome c. All data represented as Mean ± Standard Deviation (SD). N=6 for each measurement.

### Moderate to high dose of C-Mn_3_O_4_ NPs show mild to moderate inflammatory response in lungs

Sections of lung tissue from control and low dose C-Mn_3_O_4_ NP treated animals have the normal appearance of fine lace because most of the lung is composed of thin-walled alveoli (Figure 5a). The alveoli are composed of a single layer of flattened epithelial cells and pneumocytes. Between the alveoli a thin layer of connective tissue and numerous capillaries are also lined with endothelium. However, the sections from moderate and high dose groups showed signs of nonspecific interstitial pneumonia in patchy manner other than normal lung architecture. The cellular pattern consists primarily of mild to moderate chronic interstitial inflammation, containing lymphocytes and a few plasma cells, in a uniform and patchy distribution. Signs of fibrosis, honeycombing, hyaline membranes and granulomas are absent. This kind of histological features may arise from the localization of particulate matters (as evident from our biodistribution study) in lungs [25]. This features are generally reversible and do not cause long term harm to the subjects [25].

**Figure 5.**
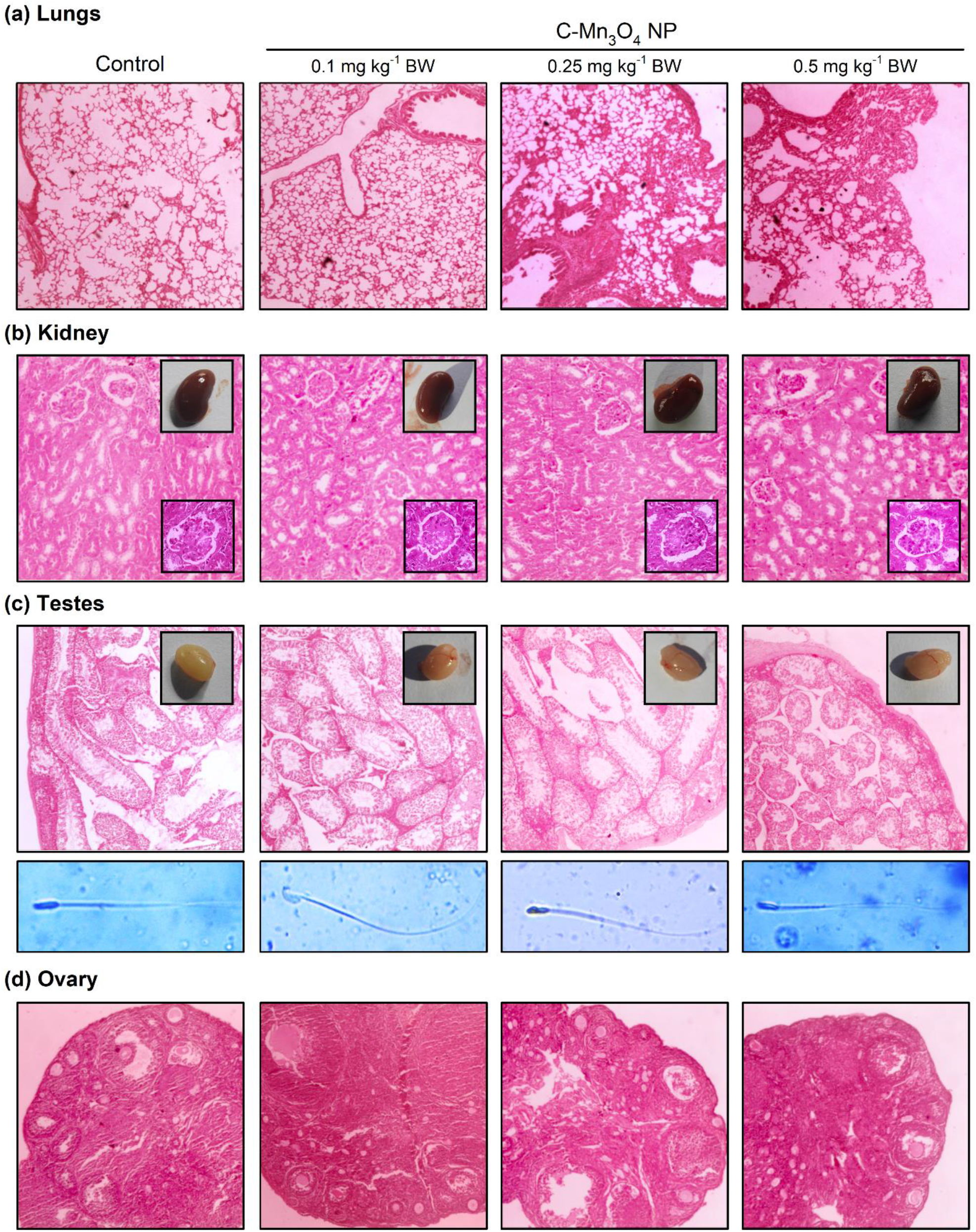
Hematoxylin and eosin stained tissue sections from different groups. (a) Lungs. (b) Kidney. Inset shows morphology of kidney and Bowman’s capsule. (c) Testes. Inset shows morphology of testes. Lower pannel shows morphology of sperm from respective groups. (d) Ovary.

### Effect of C-Mn_3_O_4_ NPs on kidney

Most of the drugs and xenobiotics pass through the kidney for excretion. But earlier studies have described that Mn cannot excrete through it. Our biodistribution results also revealed minimal deposition of Mn in the Kidney (Supplementary Figure S1). The histological analysis shows no damage and the normal structure of medulla and cortex was maintained in all four groups in spite of sexuality (Figure 5b). Consistent with these findings’ other parameters like BUN, Creatinine, Urine Volume, Urine Protein and albumin to creatinine ratio remained similar to that of the control group (Supplementary Figure S2a-S2e).

### Effect of C-Mn_3_O_4_ NPs on reproductive organs

Affinity of nanoparticles towards testes is contradictory. Several nanoparticles such as gold and magnetic nanoparticles have been reported to enter testes in small quantities, whereas carbon nanotubes behave oppositely [26]. Our biodistribution results indicate that C-Mn_3_O_4_ NPs could enter or accumulate in the testis. The average testis weight (Supplementary Figure S3a), and morphology (Figure 5c) of NP-treated mice did not show any significant difference compared to the control mice. The observed accumulation of C-Mn_3_O_4_ NPs in the testes upon chronic repeated dose administration have not shown any adverse effect on Sertoli cells, seminiferous tubules and germ cell nuclear volume (Supplementary Figure S3b-S3d). Figure 5c presents hematoxylin and eosin stained cross-section structure of the testes of control and C-Mn_3_O_4_ NP treated mice. It is well known that within the seminiferous tubules, sperm cells are generated through differentiation of spermatozoa by meiosis. Microscopic images show normal testes structure for control mice (Figure 5c). Histologic examination of the cross sections of testes from all three dose group showed no alteration even after 90 days of administration (Figure 5c). Leydig cells or spermatids were similar in all groups. The sperm counts and sperm motility showed similar trends (Supplementary Figure S3e-S3g). We further microscopically observed eosin-Y stained sperms, and found slight deformity in 0.5 mg kg^-1^ BW NP treated group (Figure 5c-lower panel). Appropriate levels of male sex hormones are crucial for spermatogenesis. However, their functions can be affected by xenobiotics or foreign substances. Some endocrine toxicants, such as polychlorinated biphenyls and inorganic heavy metals, can alter the levels of these hormones, causing injury to testes, fault in spermatogenesis and male infertility. Notably nanoparticle-rich diesel exhaust inhalation disturbs the endocrine action of the male reproductive system by increasing the plasma testosterone levels in male rats. In this study, we determined the plasma concentrations of testosterone, LH and FSH on day 90 by enzyme-linked immunosorbent assay (ELISA). The results (Supplementary Figure S3h-S3j) indicate that the NPs did not alter plasma sex hormone levels in any of the groups.

Similar to male reproductive system, no observable sign of toxicity was found in case of female reproductive system. The haematoxylin and eosin stained sections of ovary from control animals displayed normal morphology (Figure 5d). Developing follicles (primordial, primary, pre-antral and antral follicles), corpora luteum, Graafian follicles and atretic follicles were found in the cortex of the ovary which was covered by a surface epithelium (Figure 5d). The ovary of Mn_3_O_4_ NPs treated mice (at any dose) did not show any noticeable changes in histology such as formation of fibrosis or cyst. The total number of healthy follicles (all categories) remains same after chronic exposure of Mn_3_O_4_ NPs across three dose groups (Supplementary Figure S4b). The mean number of corpora lutea in mice remained same in all groups. The estrus cycle was normal with typical distribution of diestrus, estrus, proestrus and metestrus stages (Supplementary Figure S3c & S3d). The hormone levels showed no discrepancy (Supplementary Figure S3e-S3h).

### Effect of C-Mn_3_O_4_ NPs on glucose metabolism and pancreas

We further investigated the effect of C-Mn_3_O_4_ NPs on plasma glucose levels and its metabolism. Microscopic investigation of the pancreas, the major organ responsible for glucose homeostasis, both control, and NP-treated mice showed normal pancreatic tissue forming characteristic pancreatic acini with basal nuclei and amphiphilic cytoplasm (Supplementary Figure S5a). The islet of Langerhans’s displayed islet cells arranged in trabecular and acinar pattern with abundant eosinophilic cytoplasm and central small nucleus, separated by thin loose connective tissue with thin vessels. Islets have their typical spherical shape with a large number of β-cells with normal round shape distinct round nuclei. No significant change was observed across the groups for total cholesterol, triglyceride, LDL, HDL or VLDL (Supplementary Figure S5c-S5f). Plasma glucose levels (12h fasting), HbA_1c_ and serum insulin concentration remained similar to control animals (Supplementary Figure S5g & S5e).2h glucose tolerance test showed indistinctive results among the groups with identical AUC (Supplementary Figure S5i & S5j). Thus, the NPs neither affects glucose metabolism nor changes the lipid profile in the experimental regime.

## CONCLUSION

In summary, the present study intends to highlight the efficient redox buffering capability of an exclusively developed C-Mn_3_O_4_ NPs. The chronic toxicity study implies its potential safety. Furthermore, our study can serve as a template to assess other redox modulatory nanotherapeutic agents. The method is easy, semi-quantitative and sustainable for robust use. To the best of our understanding this may open a new avenue in preliminary screening and evaluation of toxicity in the redox based nanomedicine research.

## ACKNOWLEDGEMENT

The authors thank SAIF-IIT Bombay for ICP-AES measurements. MD thanks University Grants Commission (UGC), Govt. of India for Junior Research Fellowship. SKP thanks the Indian National Academy of Engineering (INAE) for the Abdul Kalam Technology Innovation National Fellowship, INAE/121/AKF. The authors thank the Department of Biotechnology (DBT, West Bengal) for the financial grant under BOOST scheme, 339/WBBDC/1P-2/2013.

## DATA AVAILABILITY

All data that support the findings of this study are available within the published article (and supplementary information files). Study related additional data will be available from SKP upon legitimate request.

## MATERIALS AND METHODS

### Quantification & characterization of *in vitro* ROS

DCFH a very popular chemical for quantification of ROS generation was prepared from DCFH-DA via de-esterification reaction at room temperature following a standardized protocol described in earlier studies [17]. ROS generated in the reaction medium reacted with DCFH and convert it to DCF which has a characteristic fluorescence emission maxima at 520 nm when excited at 488 nm. DCFH was allowed to react with increasing concentration of Ch-Mn_3_O_4_ NPs in aqueous medium and fluorescence was recorded using fluorolog, Model LFI-3751 (Horiba-Jobin Yvon, Edison, NJ) spectrofluorimeter.

### Measurement of intracellular ROS generation

After isolation RBC were incubated with C-Mn_3_O_4_ NPs (3.4 µg ml^-1^) for 3 hours. After the incubation cells were washed with PBS and treated with DCFH-DA probe for 30 minutes. At the end of this period the excess amount of DCFH-DA was thoroughly rinsed with PBS. Then cells were scraped and suspended in PBS. The cell suspensions was subjected to the measurement of DCF fluorescence intensity using fluorolog, Model LFI-3751 (Horiba-Jobin Yvon, Edison, NJ) spectrofluorimeter.

All spectroscopic experiments were performed using quartz cuvette of path length 1 cm.

### Animals

Swiss albino mice of both sex (2–3 weeks, weighing 12–15 g) were used for the current study. Four or five mice were housed together in polypropylene cage under a 12:12 light-dark cycle. During the time, food and water were accessible *ad libitum*. Mice were procured from National Institute of Nutrition, Hyderabad, India. All animal studies and experimental procedures were performed at animal house of Uluberia College, West Bengal, India (Reg. No.- 2057/GO/ReRcBi/S/19/CPCSEA) following the protocol approved by the Institutional Animal Ethics Committee (Ethical Clearance No. - 02/S/UC-IAEC/01/2019) as per standard guideline of CPCSEA, New Delhi, India.

### Biochemical tests

At the end of the experimental period, the animals were euthanized and decapitated after being fasted. Blood was collected from retro orbital plexus just before sacrifice, kept in sterile non-heparinized tubes in slanting position for 45 min and centrifuged at 3500×g for 20 min. The clear serum was obtained and used in subsequent biochemical analysis.

Biochemical evaluations were performed using commercially available kits (Autospan Liquid Gold, Span Diagnostic Ltd., India) following the protocol described by respective manufacturers. δ-aminolevulinic acid dehydratase (δ-ALAD) activity was measured using the European standardized method [27]. Complete blood count (CBC) was obtained using an automated cell counter (Medonic CA 620, Boule Diagnostics, Sweden). Hormones were estimated using commercially available ELISA kits following the protocol described by respective manufacturers.

Tissue samples were collected, homogenized in cold phosphate buffer (0.1 M; pH 7.4), and centrifuged at 10,000 r.p.m. at 4°C for 15 min. The supernatants were collected and used to determine the activity of SOD, CAT, GPx and GSH as well as the content of malondialdehyde (MDA) as marker of lipid peroxidation. Lipid peroxidation was determined in TBARS formation using a reported procedure [28, 29]. SOD (Sigma, MO, USA), CAT (Abcam, Germany), and GPx activities (Sigma, MO, USA) were estimated using commercially available test kits following protocols recommended by respective manufacturers.

### Histology

For microscopic evaluation, a conventional technique of paraffin wax sectioning and differential staining was used [30]. Organs were removed following incision, fixed in 10% neutral buffered formalin saline for 72 h, dehydrated in graduated ethanol (50–100%), cleared in xylene, and embedded in paraffin. Microtome was used to prepare ultrathin sections (4–5 μm), followed by staining with hematoxylin and eosin (H&E). Histopathological changes were examined under the microscope (Olympus BX51) equipped with a CCD based camera.

### Biodistribution

ICP-AES (ARCOS, SPECTRO Analytical Instruments GmbH, Germany) was carried out to determine the amount of Mn in blood and other organs like liver, kidney, spleen and, brain. The open acid digestion method was used for sample preparation. In brief, tissues were dried using liquid nitrogen and weighted. The freeze-dried samples were dissolved in an acid mixture that contained HNO_3_ (3 ml), H_2_SO_4_ (2 ml), and H_2_O_2_ (1 ml), heated at 120°C until only a residue remained, and then diluted with deionized water to 10 ml.

### Statistical analysis

All quantitative data are expressed as Mean±SD unless otherwise stated. One-way analysis of variance (ANOVA) followed by Tukey’s multiple comparison test was executed for comparison of different parameters between the groups using a computer program GraphPad Prism (version 8.00 for Windows), GraphPad Software (CA, USA). p<0.05 was considered significant.

## REFERENCES

1. Stone V, Donaldson K: Signs of stress. Nature Nanotechnology 2006, 1(1):23–24.

2. Wang S, Li F, Hu X, Lv M, Fan C, Ling D: Tuning the Intrinsic Nanotoxicity in Advanced Therapeutics. Advanced Therapeutics 2018, 1(5):1800059.

3. Xia T, Kovochich M, Brant J, Hotze M, Sempf J, Oberley T, Sioutas C, Yeh JI, Wiesner MR, Nel AE: Comparison of the Abilities of Ambient and Manufactured Nanoparticles To Induce Cellular Toxicity According to an Oxidative Stress Paradigm. Nano Letters 2006, 6(8):1794–1807.

4. Nel A, Xia T, Mädler L, Li N: Toxic Potential of Materials at the Nanolevel. Science 2006, 311(5761):622–627.

5. Yang B, Chen Y, Shi J: Reactive Oxygen Species (ROS)-Based Nanomedicine. Chemical Reviews 2019, 119(8):4881–4985.

6. Winterbourn CC: Reconciling the chemistry and biology of reactive oxygen species. Nature Chemical Biology 2008, 4(5):278–286.

7. Wang Z, Zhang Y, Ju E, Liu Z, Cao F, Chen Z, Ren J, Qu X: Biomimetic nanoflowers by self-assembly of nanozymes to induce intracellular oxidative damage against hypoxic tumors. Nature Communications 2018, 9(1):3334.

8. Trachootham D, Alexandre J, Huang P: Targeting cancer cells by ROS-mediated mechanisms: a radical therapeutic approach? Nature Reviews Drug Discovery 2009, 8(7):579–591.

9. Sies H, Jones DP: Reactive oxygen species (ROS) as pleiotropic physiological signalling agents. Nature Reviews Molecular Cell Biology 2020, 21(7):363–383.

10. Sims CM, Hanna SK, Heller DA, Horoszko CP, Johnson ME, Montoro Bustos AR, Reipa V, Riley KR, Nelson BC: Redox-active nanomaterials for nanomedicine applications. Nanoscale 2017, 9(40):15226–15251.

11. Nathan C, Cunningham-Bussel A: Beyond oxidative stress: an immunologist’s guide to reactive oxygen species. Nature Reviews Immunology 2013, 13(5):349–361.

12. Martinovich GG, Martinovich IV, Cherenkevich SN, Sauer H: Redox Buffer Capacity of the Cell: Theoretical and Experimental Approach. Cell Biochemistry and Biophysics 2010, 58(2):75–83.

13. Morgan B, Ezeriņa D, Amoako TNE, Riemer J, Seedorf M, Dick TP: Multiple glutathione disulfide removal pathways mediate cytosolic redox homeostasis. Nature Chemical Biology 2013, 9(2):119–125.

14. Martinovich GG, Cherenkevich SN, Sauer H: Intracellular redox state: towards quantitative description. European Biophysics Journal 2005, 34(7):937–942.

15. Giri A, Goswami N, Sasmal C, Polley N, Majumdar D, Sarkar S, Bandyopadhyay SN, Singha A, Pal SK: Unprecedented catalytic activity of Mn3O4 nanoparticles: potential lead of a sustainable therapeutic agent for hyperbilirubinemia. RSC Advances 2014, 4(10):5075–5079.

16. Adhikari A, Biswas P, Mondal S, Das M, Darbar S, Hameed AM, Alharbi A, Ahmed SA, Sankar Bhattacharya S, Pal D et al: A Smart Nanotherapeutic Agent for in vitro and in vivo Reversal of Heavy-Metal-Induced Causality: Key Information from Optical Spectroscopy. ChemMedChem 2020, 15(5):420–429.

17. Adhikari A, Polley N, Darbar S, Bagchi D, Pal SK: Citrate functionalized Mn3O4 in nanotherapy of hepatic fibrosis by oral administration. Future Science OA 2016, 2(4):FSO146.

18. Adhikari A, Mondal S, Das M, Biswas P, Pal U, Darbar S, Bhattacharya SS, Pal D, Saha-Dasgupta T, Das AK et al: Incorporation of a Biocompatible Nanozyme in Cellular Antioxidant Enzyme Cascade Reverses Huntington’s Like Disorder in Preclinical Model. Advanced Healthcare Materials 2020:2001736.

19. Horsburgh MJ, Wharton SJ, Karavolos M, Foster SJ: Manganese: elemental defence for a life with oxygen. Trends in Microbiology 2002, 10(11):496–501.

20. Takashima T, Hashimoto K, Nakamura R: Inhibition of Charge Disproportionation of MnO2 Electrocatalysts for Efficient Water Oxidation under Neutral Conditions. Journal of the American Chemical Society 2012, 134(44):18153–18156.

21. Adhikari A, Das M, Mondal S, Darbar S, Das AK, Bhattacharya SS, Pal D, Pal SK: Manganese neurotoxicity: nano-oxide compensates for ion-damage in mammals. Biomaterials Science 2019, 7(11):4491–4502.

22. Huang X, Li L, Liu T, Hao N, Liu H, Chen D, Tang F: The Shape Effect of Mesoporous Silica Nanoparticles on Biodistribution, Clearance, and Biocompatibility in Vivo. ACS Nano 2011, 5(7):5390–5399.

23. Adhikari A, Darbar S, Chatterjee T, Das M, Polley N, Bhattacharyya M, Bhattacharya S, Pal D, Pal SK: Spectroscopic Studies on Dual Role of Natural Flavonoids in Detoxification of Lead Poisoning: Bench-to-Bedside Preclinical Trial. ACS Omega 2018, 3(11):15975–15987.

24. Adhikari A, Polley N, Darbar S, Pal SK: Therapeutic Potential of Surface Functionalized Mn3O4 Nanoparticles Against Chronic Liver Diseases in Murine Model. Materials Focus 2017, 6(3):280–289.

25. Kumar V, Abbas AK, Fausto N, Aster JC: Robbins and Cotran pathologic basis of disease, professional edition e-book: Elsevier health sciences; 2014.

26. Bai Y, Zhang Y, Zhang J, Mu Q, Zhang W, Butch ER, Snyder SE, Yan B: Repeated administrations of carbon nanotubes in male mice cause reversible testis damage without affecting fertility. Nature Nanotechnology 2010, 5(9):683–689.

27. Fujita H: Measurement of δ-Aminolevulinate Dehydratase Activity. Current Protocols in Toxicology 1999, 1(1):8.6.1-8.6.11.

28. Moore K, Roberts LJ: Measurement of Lipid Peroxidation. Free Radic Res 1998, 28(6):659–671.

29. Garcia YJ, Rodríguez-Malaver AJ, Peñaloza N: Lipid Peroxidation Measurement by Thiobarbituric Acid Assay in Rat Cerebellar Slices. J Neurosci Methods 2005, 144(1):127–135.

30. Polley N, Saha S, Adhikari A, Banerjee S, Darbar S, Das S, Pal SK: Safe and Symptomatic Medicinal Use of Surface-Functionalized Mn3O4 Nanoparticles for Hyperbilirubinemia Treatment in Mice. Nanomedicine 2015, 10(15):2349–2363.

