## Supplementary Information for "Redox Buffering Capacity of Nanomaterials as an Index of ROS-based Therapeutics and Toxicity: A Preclinical Animal Study"

**Supplementary Table S1. Hematological parameters across groups.**

| Parameters | Male | | | | Female | | | |
|---|---|---|---|---|---|---|---|---|
| | **Control** | **0.1 mg $kg^{-1}$ NP** | **0.25 mg $kg^{-1}$ NP** | **0.5 mg $kg^{-1}$ NP** | **Control** | **0.1 mg $kg^{-1}$ NP** | **0.25 mg $kg^{-1}$ NP** | **0.5 mg $kg^{-1}$ NP** |
| Hb (g/dl) | 11.9±1.8 | 11.8±1.6 | 10.8±0.5 | 9.8±1.1 | 11.8±1.2 | 11.6±1.5 | 11.2±4.4 | 10.8±0.9 |
| RBC (x$10^6$ $\mu L^{-1}$) | 10.8±0.9 | 9.81±0.7 | 10.1±0.5 | 9.2±0.3 | 10.6±1.1 | 10.4±0.9 | 10.2±1.4 | 10.2±0.8 |
| RT (%) | 2.8±1.1 | 2.5±2.3 | 2.8±2.4 | 2.4±1.6 | 2.9±1.6 | 2.2±2.1 | 2.2±1.6 | 2.5±1.1 |
| HCT (%) | 34.8±1.3 | 34.8±2.2 | 32.8±2.1 | 31.8±2.1 | 35.1±3.1 | 34.9±1.6 | 33.8±2.1 | 34.1±1.4 |
| MCV ($\mu m^3$) | 37.0±2.6 | 37.4±2.4 | 36.0±1.4 | 36.1±2.3 | 37.2±1.1 | 37.4±1.9 | 37.1±1.4 | 36.0±2.4 |
| MCH (pg) | 21.1±2.4 | 21.7±1.7 | 20.1±1.7 | 21.8±2.8 | 21.4±2.6 | 21.1±1.6 | 21.4±1.6 | 20.1±2.4 |
| MCHC (%) | 41.4±2.6 | 41.2±2.1 | 40.4±1.4 | 41.7±2.4 | 41.4±1.4 | 40.6±2.1 | 41.2±1.8 | 41.4±1.6 |
| Platelets (x$10^3$ $\mu L^{-1}$) | 6.6±2.0 | 6.5±1.8 | 6.3±1.2 | 6.9±1.2 | 6.5±2.6 | 6.6±2.8 | 6.6±2.7 | 5.6±1.1 |
| WBC (x$10^5$ $\mu L^{-1}$) | 8.8±1.1 | 8.4±1.9 | 8.1±1.5 | 7.8±1.2 | 8.5±1.1 | 8.1±1.2 | 7.8±1.4 | 8.2±1.3 |
| Lymphocyte (%) | 76±6.3 | 71±5.3 | 73±5.4 | 72±3.3 | 73±3.4 | 76±3.6 | 74±1.3 | 78±2.2 |
| Neutrophil (%) | 25±6.2 | 25±5.1 | 21±5.1 | 22±4.3 | 25±6.9 | 22±6.8 | 24±5.7 | 26±5.5 |
| Monocyte (%) | 2.3±0.01 | 2.1±0.01 | 1.1±0.02 | 1.6±0.01 | 2.4±0.01 | 2.1±0.01 | 1.8±0.02 | 1.4±0.03 |
| Eosinophil (%) | 9.6±2.6 | 9.1±3.2 | 9.4±2.5 | 8.3±4.1 | 9.2±3.6 | 9.1±1.4 | 9.4±2.9 | 9.2±1.2 |
| Basophil (%) | 1.2±0.05 | 1.1±0.04 | 1.2±0.02 | 1.5±0.02 | 1.2±0.04 | 1.1±0.02 | 1.1±0.01 | 1.3±0.04 |
| PT (sec) | 7.3±2.2 | 7.2±2.1 | 8.5±3..9 | 9.1±3.8 | 7.1±4.1 | 7.4±2.6 | 8.2±3.1 | 9.3±3.1 |
| APTT (sec) | 16.9±4.3 | 16.5±2.8 | 17.1±4.1 | 18.7±2.6 | 16.1±2.5 | 15.2±2.6 | 16.4±3.2 | 19.6±4.2 |

Data are expressed as mean ± standard deviation (N=06).

Abbreviations: Hb: Hemoglobin; RBC: Red Blood corpuscle; RT: Reticulocyte; HCT: Hematocrit; MCV: Mean Corpuscular Volume; MCH: Mean Corpuscular Hemoglobin; MCHC: Mean Corpuscular Hemoglobin Concentration; WBC: White Blood Corpuscle; PT: Prothrombin Time; APTT: Activated Partial Thromboplastin Time.

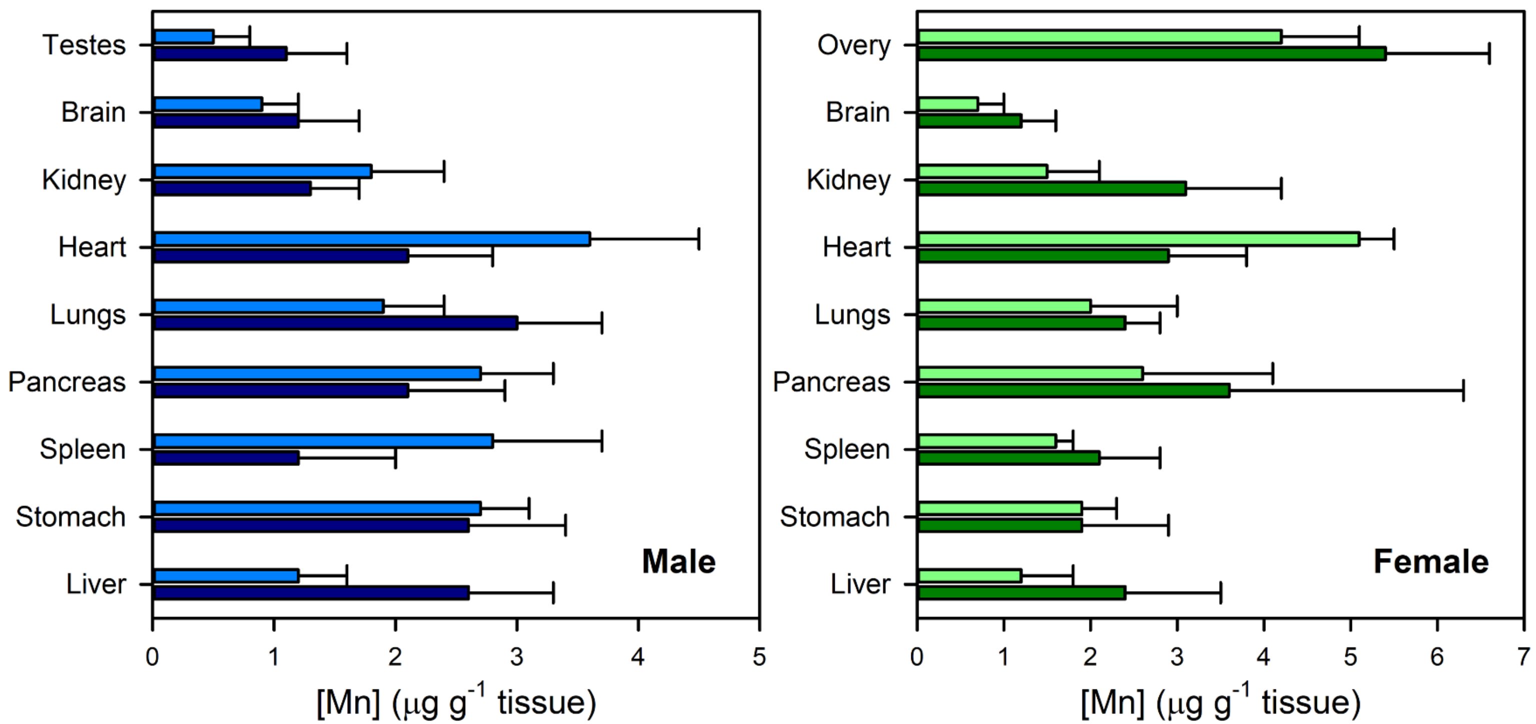


**Supplementary Figure S1. Biodistribution of C-$Mn_3O_4$ NPs in various organs after 90 days of oral exposure.**

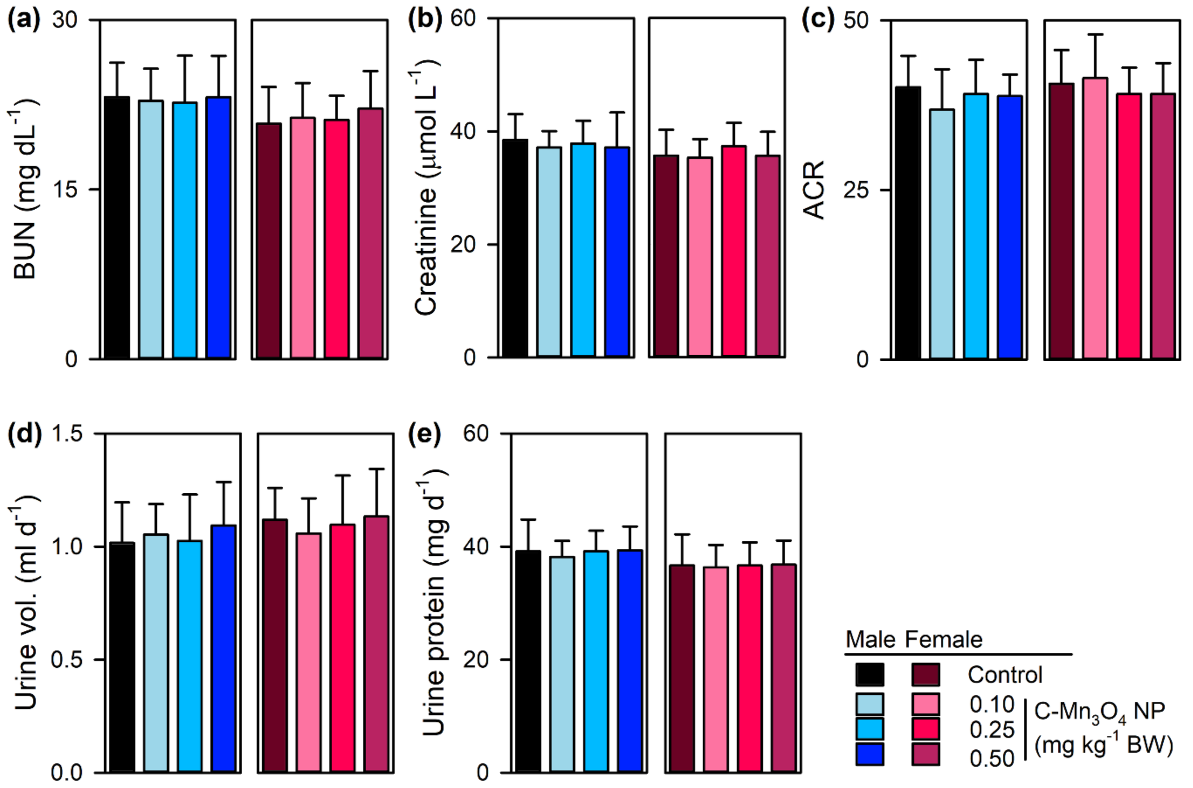


**Supplementary Figure S1. Effect of C-$Mn_3O_4$ NPs on renal system.** (a-e) Effect of C-$Mn_3O_4$ NPs on kidney antioxidant enzyme functions.

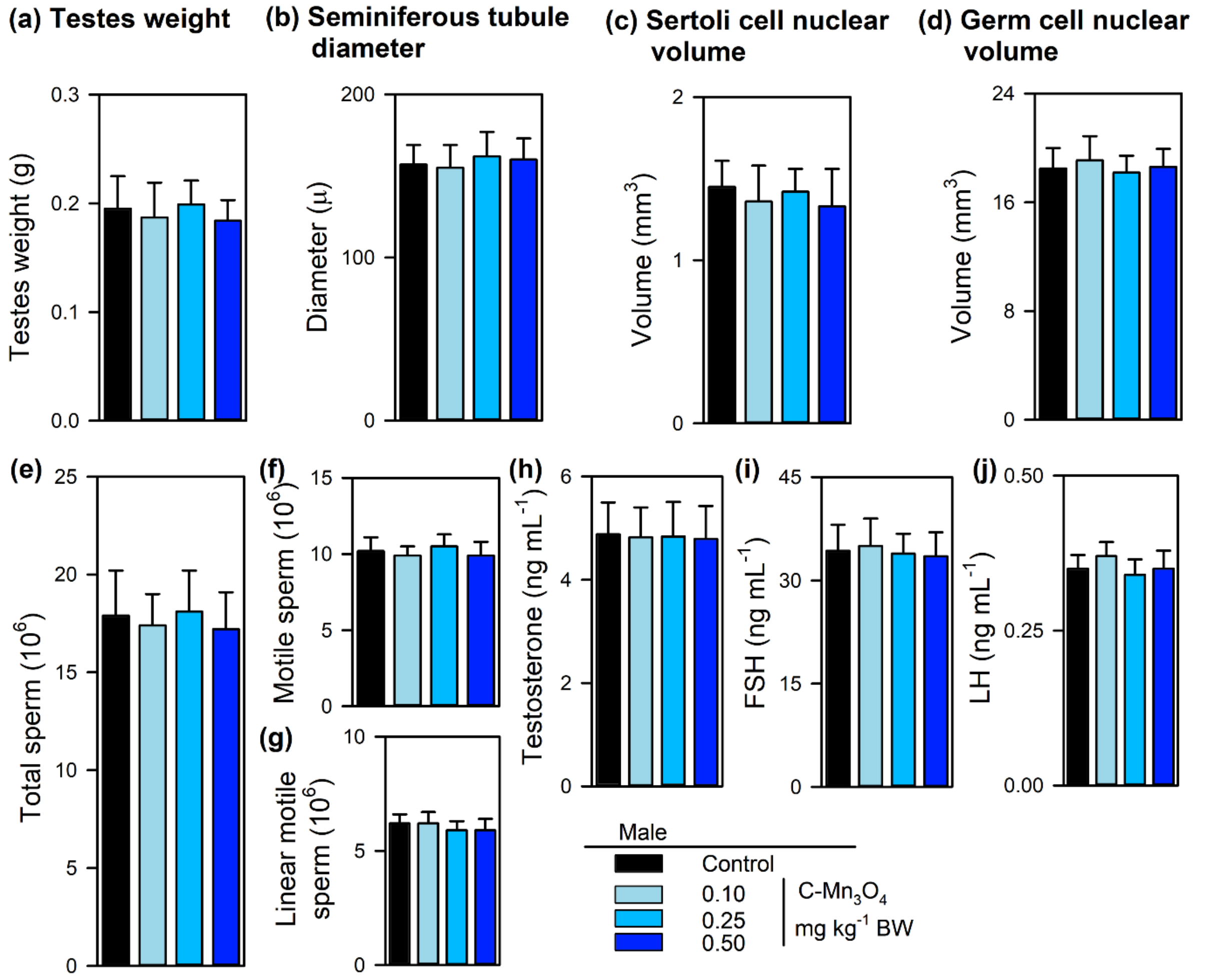


**Supplementary Figure S2. Effect of C-$Mn_3O_4$ NPs on male reproductive system.** (a) Weight of testes. (b-g) The quantity of different types of spermatogonia as observed in histology. (e-g) Sperm counts and sperm viability has no difference in between groups. Micrographs of single sperm (stained with Eosin Y) depicts the same. (h-j) Male reproductive hormones remained unchanged across the groups.

All data represented as Mean ± Standard Deviation (SD). N=6 for each measurement.

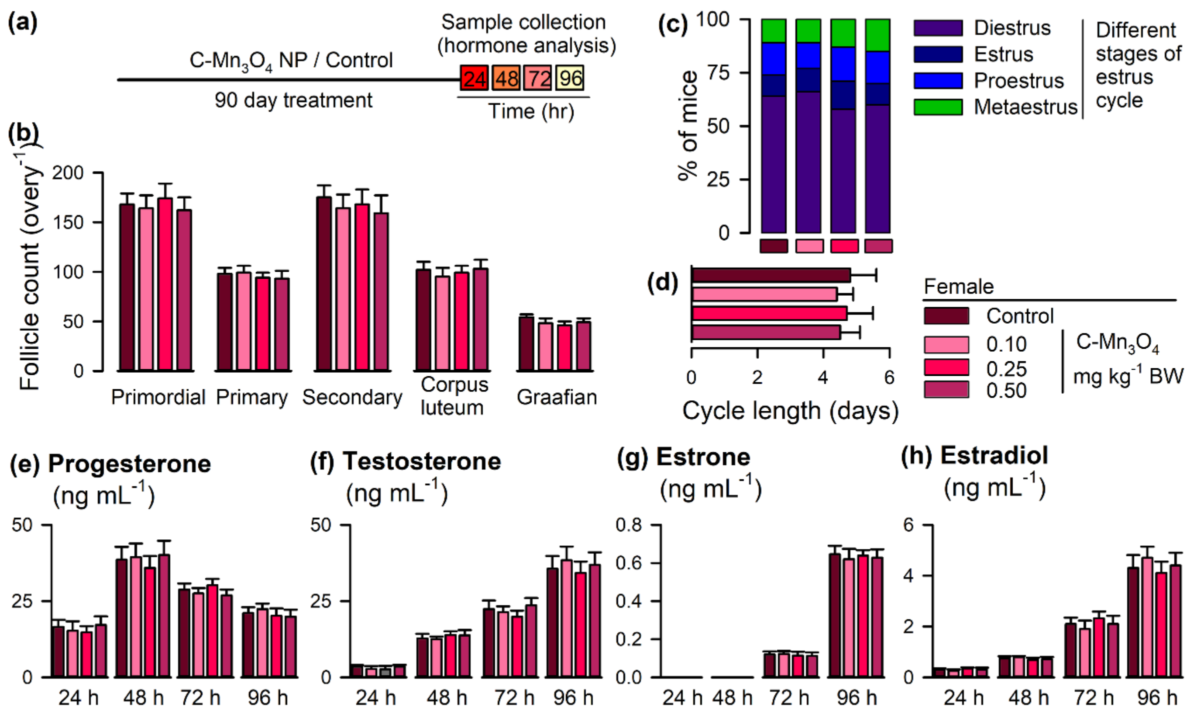


**Supplementary Figure S3. Effect of C-$Mn_3O_4$ NPs on female reproductive system.** (a) Collection protocol. (b) Differential counts of the follicles as observed in histological sections. (c) Distribution of estrus cycle across the groups. (d) Cycle length. (e-h) Changes in female reproductive hormones due to chronic C-$Mn_3O_4$ NPs exposure.

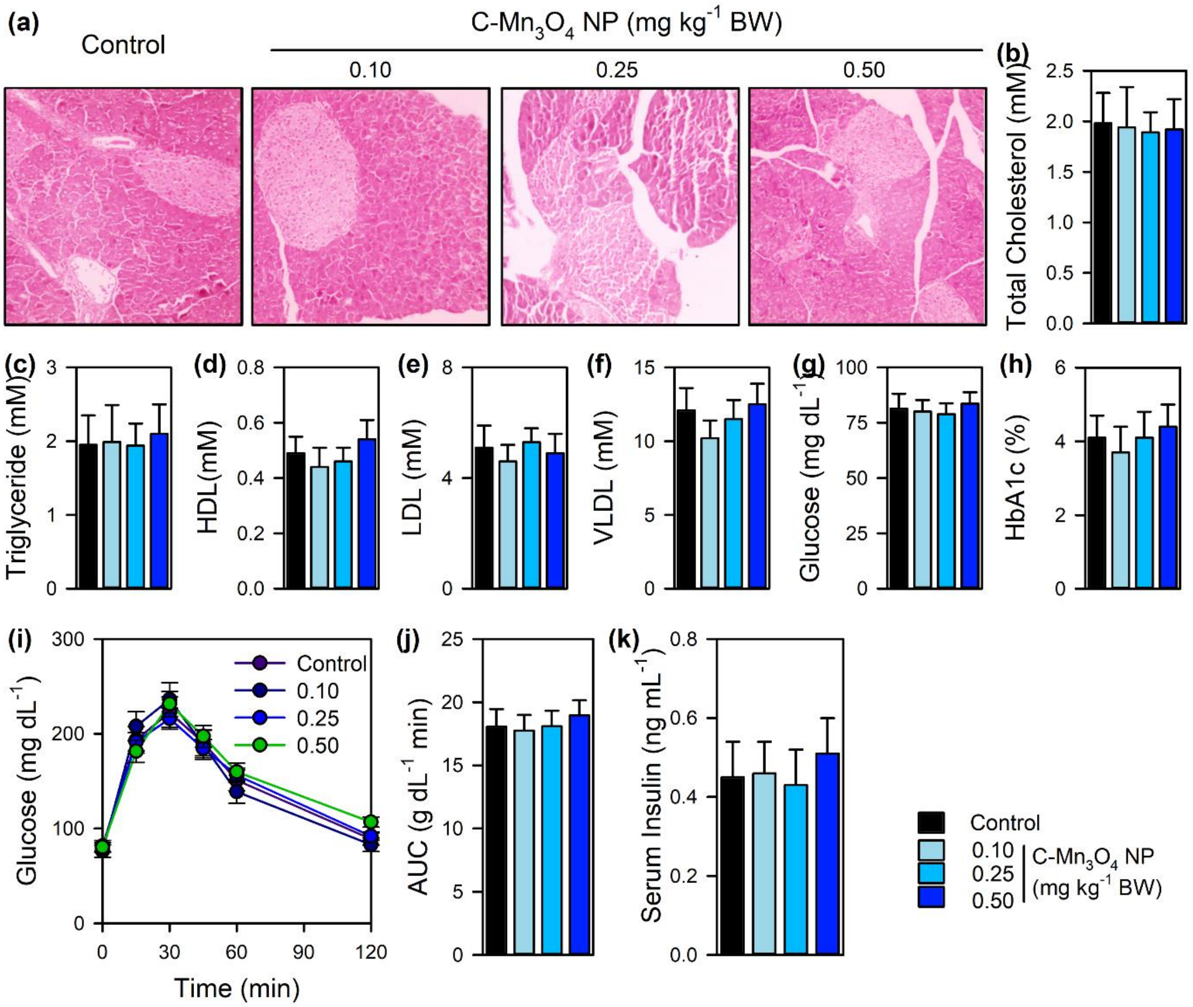


**Supplementary Figure S4. Effect of the C-$Mn_3O_4$ NPs on glucose metabolism.** (a) H&E stained sections of pancreas. (b) Glucose uptake test. (c) AUC and other glucose metabolism parameters. (d) Lipid profile of the BALB/c mice treated with C-$Mn_3O_4$ NPs for 90 days.
